# Cancer cells induce hepatocytes apoptosis in co-opted colorectal cancer liver metastatic lesions

**DOI:** 10.1101/2021.02.11.429243

**Authors:** Miran Rada, Migmar Tsamchoe, Audrey Kapelanski-Lamoureux, Jessica Bloom, Stephanie Petrillo, Diane H Kim, Anthoula Lazaris, Peter Metrakos

## Abstract

Vessel co-option in colorectal cancer liver metastases (CRCLM) has been recognized as one of the mechanistic pathways that contribute to resistance against anti-angiogenic therapy. In vessel co-opted CRCLM lesions, the cancer cells are highly motile that move toward and along the pre-existing sinusoidal vessels and hijack them to gain access to nutrient. The movement of cancer cells is accompanied by replacement of the hepatocytes. However, the molecular mechanisms by which this replacement occurs are unclear yet. To examine the involvement of apoptosis in hepatocytes replacement by cancer cells in co-opted lesions, we conducted immunohistochemical staining for chemonaïve CRCLM specimens using pro-apoptotic markers antibody, such as cleaved caspase-3 and cleaved poly (ADP-ribose) polymerase-1 (PARP-1). The results suggested overexpression of pro-apoptotic markers in liver parenchyma of co-opted lesions compared to angiogenic lesions, specifically the hepatocytes that are in close proximity to the cancer cells. Importantly, co-culturing hepatocytes with colorectal cancer cells induced overexpression of pro-apoptotic markers in the hepatocytes. Altogether, these results propose that cancer cells could exploit apoptosis to replace the hepatocytes and establish vessel co-option in CRCLM.

## 1. Introduction

Colorectal cancer (CRC) is the third most commonly diagnosed cancer and the second deadliest malignancy worldwide [1]. Metastases accounts for most of CRC linked deaths, which 25%-30% of patients will develop liver metastases [2]. Surgical resection is currently the most effective treatment approach for CRCLM patients that results in 5-year survival rates of up to 50%. However, only 20% of CRCLM patients are suitable for upfront surgery and the remains are left with palliative options [3,4].

Pathologists have noted two major histological heterogeneity within CRCLM including desmoplastic histological growth pattern (DHGP) and replacement histological growth pattern (RHGP) [5–8]. The DHGP lesions are characterized by desmoplastic stroma separating liver parenchyma from tumour, whereas in RHGP lesions the tumour cells infiltrate through liver parenchyma and replacing the hepatocytes in a random fashion and following the sinusoidal structure [6,8]. Importantly, the cancer cells in DHGP lesions obtain blood supply through angiogenesis, while their counterparts in RHGP lesions co-opt the pre-existing blood vessel to gain access to nutrients [6,8–10].

To co-opt the sinusoids for blood supply in RHGP lesions, the cancer cells require high levels of motility in order to infiltrate through hepatocytes [6,9]. The infiltration of cancer cells is accompanied by replacement of the hepatocytes [11]. However, the mechanisms by which cancer cells replace hepatocytes are largely unknown. Previous studies hypothesized that various mechanisms may be involved in the hepatocytes displacement by cancer cells, such as apoptosis, necrosis or autophagy [11].

Apoptosis is a highly organized process, which cells are fragmented into small membrane-bound apoptotic bodies with cleaved DNA and proteolytic fragments that subsequently cleared by phagocytosis [12]. Cleavage of poly (ADP-ribose) polymerase-1 (PARP-1) by caspases (e.g. caspase-3) is considered as a hallmark of apoptosis [13,14]. Apoptosis play a vital role in cancer [15–18]. Evasion of apoptosis is one of the hallmarks of cancer resistance to therapeutic agents [16,19]. While some investigations have been conducted to analyse the expression levels of apoptotic genes in non-angiogenic cancers, the role of apoptosis in vessel co-option is yet not well understood. In this context, Hu et al. [20] has reported higher levels of pro-apoptotic genes including FOS, FAH, and PRODH in non-angiogenic non-small-cell lung cancer lesions comparing to their angiogenic counterparts [20,21]. Conversely, the anti-apoptotic genes including ANXA7 and SOD1 were highly expressed in angiogenic lesions [20].

In the current study, we report that apoptosis seems to play an essential role in the development of co-opted CRCLM lesions, which cancer cells in tumour–liver interface promote pro-apoptotic environment to gradually replace the hepatocytes and expand to facilitate the co-option of the intervening pre-existing sinusoidal blood vessels.

## 2. Results

### 2.1. Pro-apoptotic markers upregulated in the hepatocytes that are in close proximity to cancer cells

In the co-opted CRCLM lesions, cancer cells form cell plates that are in continuity with liver cell plates, which allows the cancer cells to replace the hepatocytes within the liver cell plates to hijack the sinusoidal blood vessels at the tumour–liver interface (Figure 1a) [5]. However, the mechanisms underlying hepatocytes replacement by cancer cells are still unclear. Therefore, we questioned whether the cancer cells in the interface areas of co-opted lesions induce apoptosis in adjacent hepatocytes to facilitate their replacement. To address this question, we performed immunohistochemical staining on a total of ten chemonaïve CRCLM specimens (n=5 DHGP and n=5 RHGP) using anti cleaved casapase-3, a biomarker for apoptosis [16,19,22,23]. As shown in Figure 1b and 1c, we observed significant increase of cleaved caspase-3 expression in hepatocytes that are in close proximity to cancer cells in co-opted RHGP lesions. Conversely, negative staining of cleaved caspase-3 was observed in the adjacent hepatocytes of angiogenic DHGP lesions. Similar results were noticed when we stained CRCLM lesions with cleaved PARP-1 antibody (Figure 1d and 1e), pro-apoptotic marker [16,19,22,23]. Importantly, we also noticed high levels of positively stained immune cells in the replacement lesions for both cleaved caspase-3 and cleaved PRAP-1. Since higher levels of neutrophils in normal liver parenchyma have been observed in co-opted CRCLM lesions comparing to angiogenic lesions [10], we hypothesized that the apoptotic immune cells might be neutrophils. Indeed, our immunofluorescence staining confirmed that the majority of the apoptotic immune cells are neutrophils (Figure S1). Further investigations are required to determine whether cancer cells are responsible for inducing apoptosis in the immune cells of co-opted CRCLM lesions, as well as identify their correlation with vessel co-option development.

**Figure 1.**
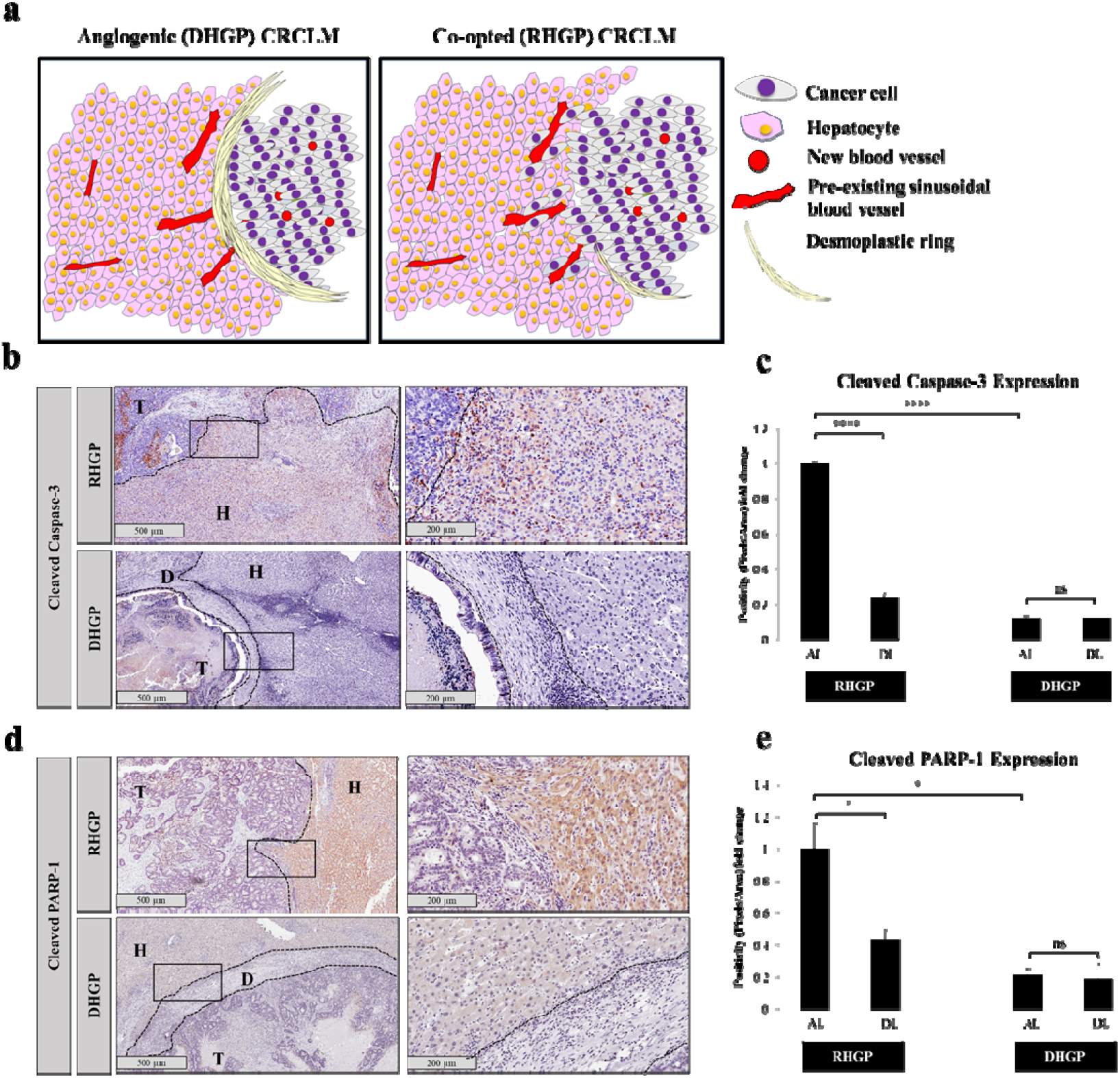
Co-opted CRCLM lesions overexpress pro-apoptotic markers. **a.** Schematic figure represent angiogenic desmoplastic (DHGP) versus co-opted replacement CRCLM lesions. **b.** Immunohistochemistry staining of chemonaïve CRCLM lesions with cleaved caspase-3 antibody. **c.** Represents quantification of cleaved caspase-3 positive staining in RHGP (n=5) and DHGP (n=5) was performed using Aperio software and positive pixels were quantified and are expressed as percent positive pixels (mean + SD). AL=Adjacent liver, DL=Distal liver. **d.** Immunohistochemistry staining of chemonaïve CRCLM lesions with cleaved PARP-1 antibody. **e.** Shows quantification of cleaved PARP-1 positive staining in RHGP (n=5) and DHGP (n=5) was performed using Aperio software and positive pixels were quantified and are expressed as percent positive pixels (mean + SD). T:Tumour; H:Hepatocytes; D:Desmoplastic ring. AL=Adjacent liver, DL=Distal liver. Error bar, SD; ns: not significant; *p < 0.05; ****p < 0.0001 (unpaired t-test).

To further confirm that the staining of both cleaved caspase-3 and cleaved PARP-1 are primarily localized in the hepatocytes, we conducted co-immunofluorescent staining for chemonaïve CRCLM lesions using hepatocyte specific antigen (HSA) and anti cleaved caspase-3 or cleaved PARP-1 antibodies. The results showed co-localization of cleaved caspase-3 and HSA (Figure 2A), as well as cleaved PARP-1 and HSA (Figure 2B). Taken together, these data indicate that the adjacent hepatocytes in the co-opted CRCLM lesions are more apoptotic comparing to their counterparts in the angiogenic lesions.

**Figure 2.**
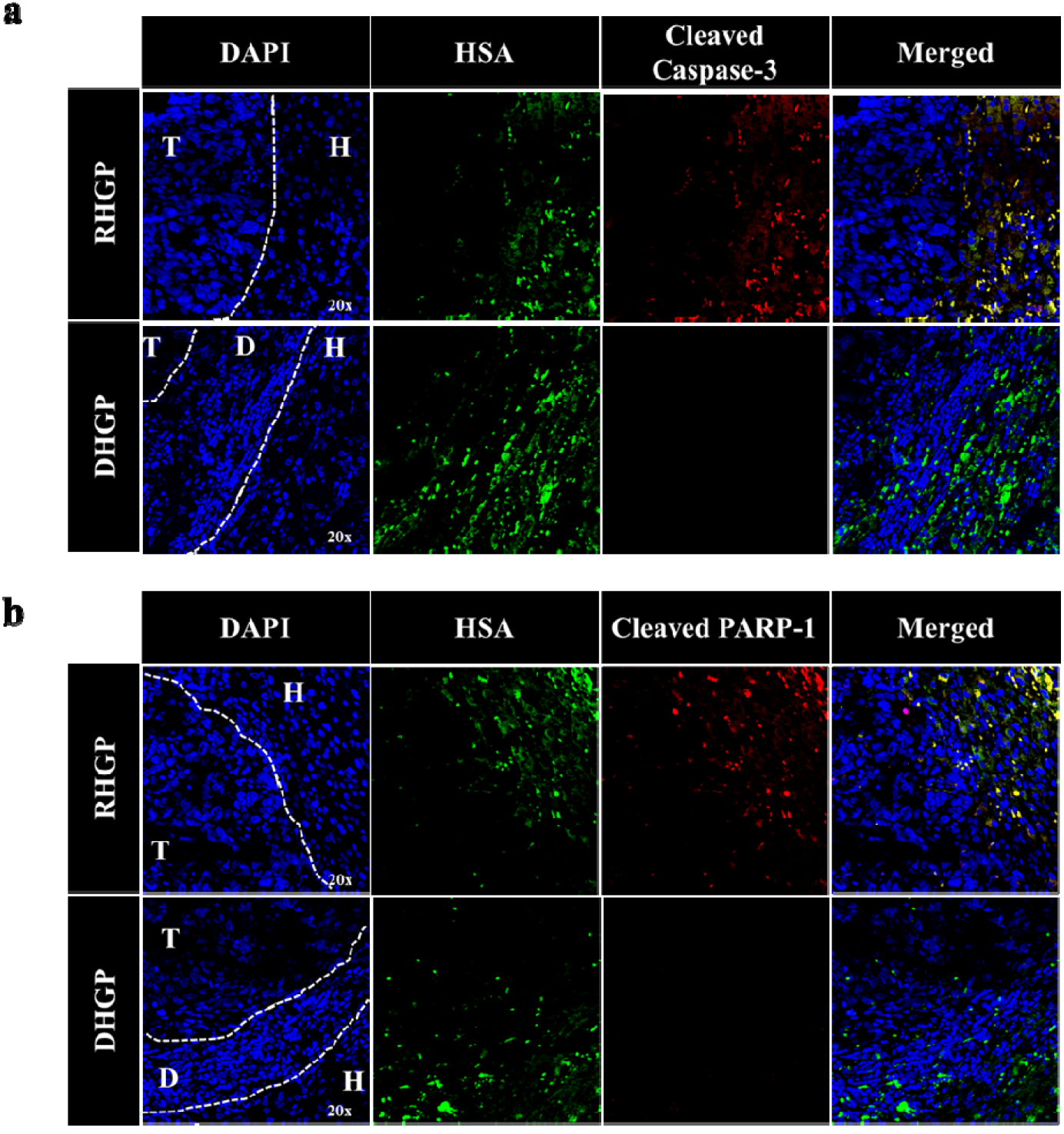
Pro-apoptotic markers are mainly localized in the hepatocytes of co-opted CRCLM lesions. **a-b.** Immunofluorescence staining of chemonaïve CRCLM lesions showing HSA (green) and cleaved caspase-3 (red) or cleaved PARP-1 (red). T:Tumour; H:Hepatocytes; D:Desmoplastic ring.

### 2.2. Cancer cells promote apoptosis in hepatocytes

To examine the association between cancer cells and hepatocytes apoptosis, we analysed the expression of pro-apoptotic markers (cleaved caspase-3 and cleaved PARP-1) in IHH hepatocytes upon their contact with colorectal cancer cells (LS174, SW620 or HT29) using insert co-culturing approach (Figure 3a). IHH cells are immortalized human hepatocytes that retained several differentiated features of normal hepatocytes [24–26]. Interestingly, the expression of both cleaved caspase-3 and cleaved PARP-1 in hepatocytes was incited upon their co-culturing with cancer cells (Figure 3b). These data indicate that apoptosis in hepatocytes could be stimulated by cancer cells in co-opted CRCLM lesions. This process is mediated by various mediators that secreted by cancer cells (Figure 4), which further studies are needed to identify these proteins.

**Figure 3.**
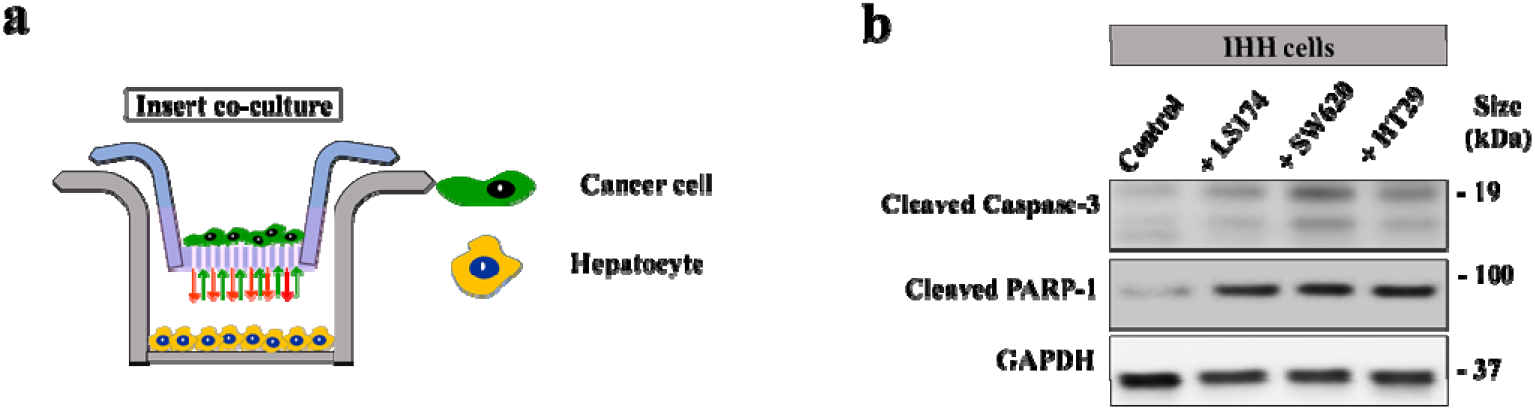
Colorectal cancer cells promote apoptosis in hepatocytes in vitro. **a.** Schematic of experimental strategy to co-culture colorectal cancer cells and IHH hepatocytes. **b.** Western blot of cleaved caspase-3 and cleaved PARP-1 in hepatocytes (IHH cells) co-cultured with different cancer cell lines.

**Figure 4.**
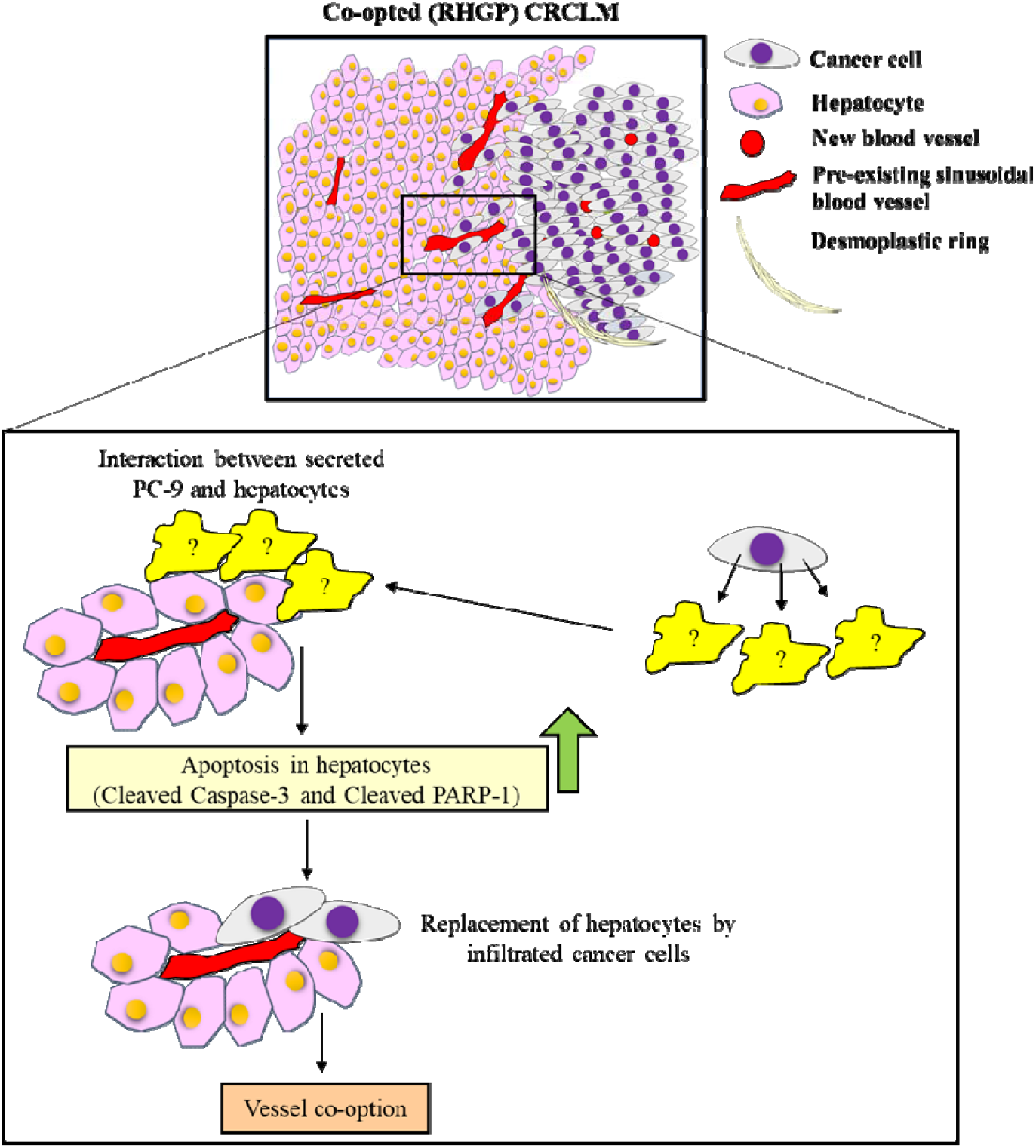
cancer cells enhance hepatocytes apoptosis through unknown mediators. Schematic summary showing the role of apoptosis in the development of co-opted RHGP lesions in CRCLM.

## 3. Discussion

The histopathological growth patterns (HGPs) in CRCLM play a vital role in tumour response to anti-angiogenic therapy [6,10]. However, the factors that dictate the development of various HGPs are yet unclear, as well as the stages of their development. In CRCLM, it has been theorized that the co-opted replacement lesions develop in four various fluid phases: a. elevation in cancer cells motility that allows the cancer cells to infiltrate through liver parenchyma, b. replacement of hepatocytes by cancer cells, c. interaction between cancer cells and pre-existing vasculature, d. recruitment of immune components that facilitate the establishment of vessel co-opting micro-environment [11]. Thus, hepatocytes replacement by cancer cells appears to be pivotal in the development of co-opted CRCLM lesions. However, the approaches of hepatocytes displacement by cancer cells are poorly understood. Herein, we observed significantly higher expression of pro-apoptotic markers in the hepatocytes of co-opted replacement lesions in comparison to angiogenic desmoplastic lesions. We therefore report hepatocytes apoptosis as one of the existing displacement mechanisms that could be used by cancer cells to displace the hepatocytes and establish vessel co-option in CRCLM.

The expression of pro-apoptotic and anti-apoptotic markers in non-angiogenic tumours or anti-angiogenic resistant cancer cells has been investigated previously [20,27–29]. In contrary to our study, most of the studies were focused on the apoptosis in cancer cells and neglected tumour microenvironment and stromal cells. In this context, Hu et al. [20] have reported lower expression levels of pro-apoptotic genes in nonangiogenic non-small-cell lung cancer cells, such as FOS, FAH, and PRODH. This finding is consistent with our results that showed negative staining of cancer cells for pro-apoptotic markers (cleaved caspase-3 and cleaved PRAP-1). On the other hand, Valient et al. [28] observed overexpression of serpin B2 (SERPINB2) in co-opted metastatic cancer cells in brain, which plays a key role in apoptosis suppression [28,30]. Therefore, it would be interesting to determine the function of SERPINB2 in apoptosis in co-opted CRCLM lesions.

Our data suggest high number of neutrophils in co-opted lesions that expressed pro-apoptotic biomarker (cleaved PARP-1) (Supplementary Figure S1). Interestingly, increasing neutrophils death has been found to promote cancer cell proliferation, adhesion, migration, invasion, and thereby tumour metastasis [31]. The role of apoptotic neutrophils in the development of co-opted CRCLM lesions is beyond the scope of this article and need to be further elucidated.

## 5. Conclusions

Overall, our data suggest apoptosis as one of the mechanisms that employed by cancer cells to replace the hepatocyte and develop vessel co-option in CRCLM. The next step is to identify the secreted proteins that mediate the induction of hepatocytes apoptosis by cancer cells. Moreover, it would be interesting to identify whether blocking hepatocytes apoptosis will be efficient to attenuate vessel co-option development or convert co-opted lesions to angiogenic in vivo.

## 4. Materials and methods

### 4.1. Patient samples

The study was conducted in accordance with the guidelines approved by McGill University Health Centre Institutional Review Board (IRB). Informed consent was obtained from all patients through the McGill University Health Centre (MUHC) Liver Disease Biobank. Surgical specimens were procured and released to the Biobank immediately after the pathologist’s confirmation of carcinoma and surgical margins.

### 4.2. Cells Co-culture and treatment

Human colorectal cancer (HT29, LS174, SW620) cell lines were a gift, kindly supplied by Dr Alex Alex Gregorieff (Cancer Research Program, McGill University). IHH cells were a generous gift from Dr Nabil G. Seidah at Montreal Clinical Research Institute (IRCM). The cells were cultured in in DMEM (Wisent Inc., #319-005-CL) supplemented with 10% FBS (Wisent Inc., #085-150) and 1× penicillin/streptomycin (Wisent Inc., 450-201-EL). All cells were cultured at 37 °C with 5% CO2.

Co-culturing was conducted using 6-well inserts (Falcon, #353090) and companion plates (Falcon #353502). The cells were seeded in DMEM (Wisent Inc., #319-005-CL) supplemented with 10% FBS (Wisent Inc., #085-150) and 1× penicillin/streptomycin (Wisent Inc., 450-201-EL) overnight. Next day, the condition media was aspirated, and the cells were washed with PBS (Wisent Inc., # 311-010-CL) twice. Then, serum-free DMEM media was added and the cells were incubated for 48 h at 37 °C.

### 4.3. Immunoblotting

To prepare lysate from the cultured cells, the medium was removed, and cells were washed once with 1x PBS, trypsinized, collected and kept on ice. Cells were resuspended in RIPA buffer (Sigma Aldrich, #R0278) supplemented with protease inhibitor (Sigma Aldrich, #4693124001), and ruptured by passing through a syringe 10 times and centrifuged for 10 minutes at 5000 rpm. The supernatant was transferred into 1.5mL microcentrifuge tubes, and protein concentrations were determined using BCA Protein Assay Kit (Thermo Scientific, #23225). 5–10 μg of total protein per sample were subjected to 10-12% SDS-PAGE and transferred to Immobilon-E membranes (Millipore, #IEVH85R). The blots were developed using Pierce ECL Western Blotting Substrate (Thermo Scientific, #32106) and imaged with ImageQuant LAS4000 (GE Healthcare BioScience), as previously described [9].

The following primary antibodies were used: GAPDH 1:2000 (Abcam, # ab9485), cleaved caspase-3 1:500 (Cell signaling, # 9664S) and cleaved PARP-1 1:500 (Cell signaling, # 5625S).

### 4.4. Immunohistochemical staining

Formalin-fixed paraffin-embedded (FFPE) CRCLM resected blocks were used for this study. Serial sections 4 mm thick were cut from each FFPE block, mounted on charged glass slides (Fisher Scientific, #12-550-15) and baked at 37°C overnight. Prior staining, the slides were baked at 60°C for 1 hr as well. Hematoxylin and eosin (H&E)-stained sections were prepared from all cases for an initial histopathological assessment. The sections were deparaffinized with xylene (Leica, #3803665) followed by hydration with graded concentrations of ethanol (Comalc, #P016EAAN) and then with distilled water. Samples were subjected to antigen retrieval (pH = 6.0 was used for cleaved caspase-3, while pH = 9.0 was used for cleaved PARP-1) followed by washing with PBS and incubation in hydrogen peroxide (Dako, #S2003) to inhibit endogenous peroxidase. The tissue sections were blocked with 1 % goat serum and incubated with the indicated primary antibody in 1 % goat serum overnight at 4°C. After washing, the sections were incubated with secondary antibody (Dako, Anti-Mouse: #K4001; Anti-Rabbit: #K4003) for 1 h at room temperature and positive signals were visualized with the diaminobenzidine (DAB) substrate (Dako, #K3468).

The following primary antibodies were used: cleaved caspase-3 1:100 (Cell signaling, # 9664S) and cleaved PARP-1 1:50 (Cell signaling, # 5625S).

All slides were scanned at 20× magnification using the Aperio AT Turbo system. Images were viewed using the Aperio ImageScope ver.11.2.0.780 software program for scoring analysis and assessment of signals. The positivity [Total number of positive pixels divided by total number of pixels: (NTotal – Nn)/(NTotal)] was assessed with an Aperio ScanScope (Aperio Technologies Inc., Vista, CA) [8–10].

### 4.5. Immunofluorescence staining

Formalin-fixed paraffin-embedded (FFPE) human CRCLM resected blocks were deparaffinized with xylene followed by hydration with graded concentrations of ethanol and then with distilled water. Samples were subjected to antigen retrieval antigen retrieval (pH = 6.0 for cleaved caspase-3 and pH = 9.0 for cleaved PARP-1) followed by washing with PBS and incubation in hydrogen peroxide (Dako, #S2003) to inhibit endogenous peroxidase. The tissue sections were blocked with 1 % goat serum and incubated with the indicated primary antibody in 1 % goat serum overnight at 4°C. After washing, the sections were incubated with secondary antibody 1:1000 (Alexa Flour 594 goat anti-rabbit IgG and Alexa Flour 488 goat anti-mouse IgG (Invitrogen #A11037 and #A10680 respectively)) for 1 h at room temperature followed by washing thrice. The sections were incubated with 4′,6-Diamidino-2-Phenylindole, Dihydrochloride DAPI 1:1000 (Thermo Fisher Scientific, D1306) in PBS for 10 minutes at room temperature. Prior to mounting under cover glasses, 1-2 drops of ProLong Gold Antifade Mountant (Thermo Fisher Scientific, P36934) were added to each section.

The following primary antibodies were used: HSA 1:300 (Santa Cruz, #SC5893), leaved caspase-3 1:100 (Cell signaling, # 9664S), cleaved PARP-1 1:50 (Cell signaling, # 5625S), elastase 1:250 (R&D Systems, #MAB91671).

### 4.6. Statistical analysis

Statistical analysis was performed with a two-tailed Student’s t-test using Excel software. Unpaired t-test was applied for compare means of two groups. P-values of <0.05 were considered to be significant.

## Author contributions

Conceptualization, M.R., A.L. and P.M.; methodology, M.R., M.T., A.K., J.B., S.P., S.T., D.H.K., P.Y.; resources, P.M., data curation, M.R.; writing _ original draft preparation, M.R.; writing – review and editing, M.R., A.L., P.M., funding acquisition, P.M. All authors have read and agreed to the published version of the manuscript.

## Acknowledgments

The authors would like to acknowledge the support that provided by Dana Massaro and Ken Verdoni Liver Metastases Research Fellowship. We thank RI-MUHC Liver Disease Biobank for providing CRCLM slides. We also thank all those patients who participated in this study and whose samples were pivotal to this research.

## Figure legends

**Supplementary Figure S1.**
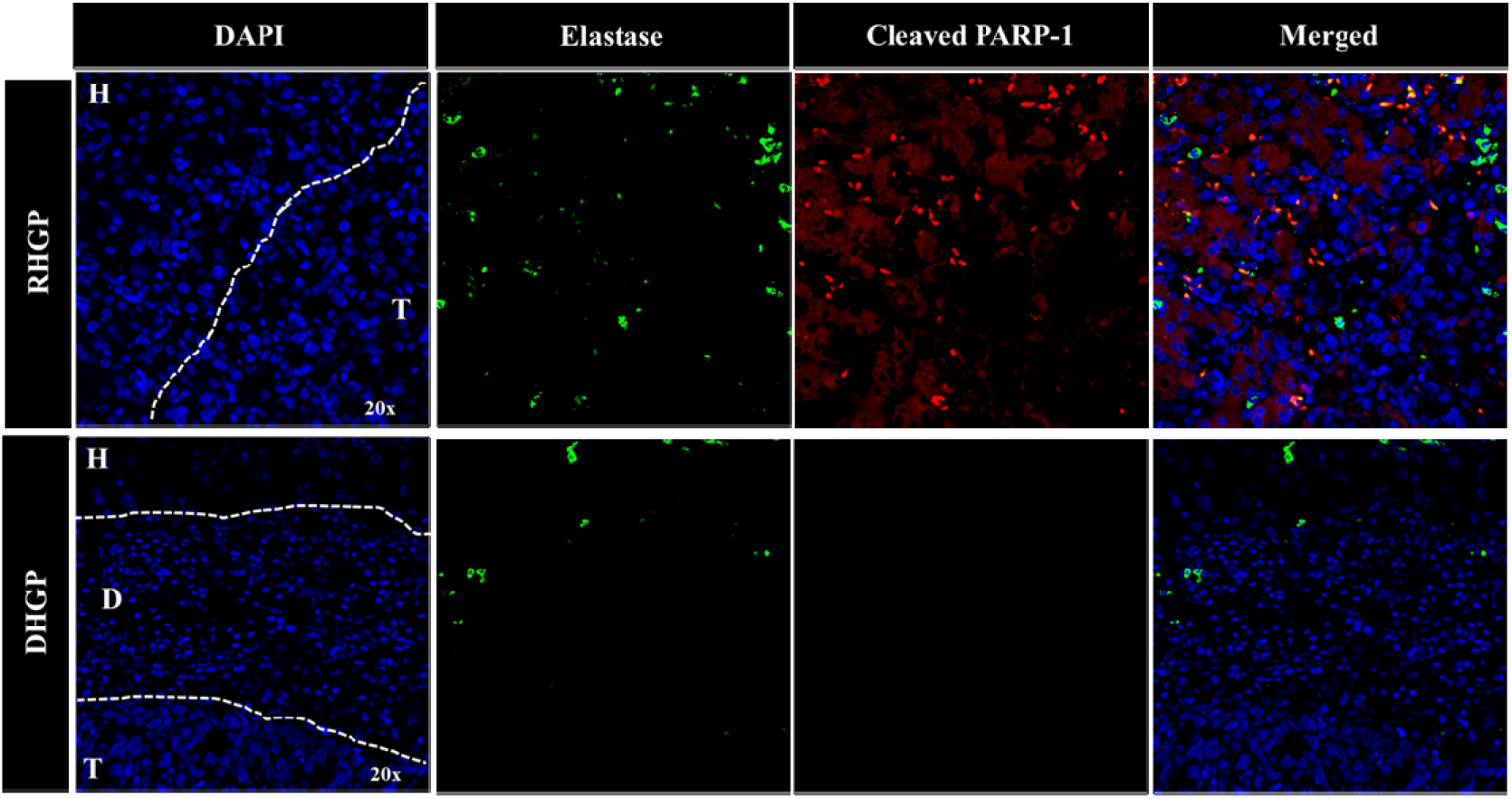
The majority of neutrophils in co-opted CRCLM lesions are apoptotic. Immunofluorescence staining of chemonaïve CRCLM lesions showing neutrophil biomarker elastase (green) and apoptosis biomarker cleaved PARP-1 (red). T:Tumour; H:Hepatocytes; D:Desmoplastic ring.

## References

[1] Bray F, Ferlay J, Soerjomataram I. Global Cancer Statistics 2018☐: GLOBOCAN Estimates of Incidence and Mortality Worldwide for 36 Cancers in 185 Countries. CA Cancer J Clin 2018;68:394–424. https://doi.org/10.3322/caac.21492.

[2] Bingham G, Shetye A, Suresh R, Mirnezami R. Impact of primary tumour location on colorectal liver metastases: A systematic review. World J Clin Oncol 2020;11:294–307. https://doi.org/10.5306/wjco.v11.i5.294.

[3] Fusai G, Davidson BR. Strategies to Increase the Resectability of Liver Metastases from Colorectal Cancer. Dig Surg 2003;20:481–96. https://doi.org/10.1159/000073535.

[4] Zarour LR, Anand S, Billingsley KG, Bisson WH, Cercek A, Clarke MF, et al. Colorectal Cancer Liver Metastasis☐: Evolving Paradigms and. Cell Mol Gastroenterol Hepatol 2017;3:163–73. https://doi.org/10.1016/j.jcmgh.2017.01.006.

[5] Dam P V, Van Der Stok EP, Teuwen LA, Van Den Eynden GG, Illemann M, Frentzas S, et al. International consensus guidelines for scoring the histopathological growth patterns of liver metastasis. Br J Cancer 2017;117:1427–41. https://doi.org/10.1038/bjc.2017.334.

[6] Frentzas S, Simoneau E, Bridgeman VL, Vermeulen PB, Foo S, Kostaras E, et al. Vessel co-option mediates resistance to anti-angiogenic therapy in liver metastases. Nat Med 2016;22:1294–302. https://doi.org/10.1038/nm.4197.

[7] Eynden GG Van den, Bird NC, Majeed AW, Van Laere S, Dirix LY, Vermeulen PB. The histological growth pattern of colorectal cancer liver metastases has prognostic value. Clin Exp Metastasis 2012;29:541–9. https://doi.org/10.1007/s10585-012-9469-1.

[8] Lazaris A, Amri A, Petrillo SK, Zoroquiain P, Ibrahim N, Salman A, et al. Vascularization of colorectal carcinoma liver metastasis: insight into stratification of patients for anti-angiogenic therapies. J Pathol Clin Res 2018;4:184–92. https://doi.org/10.1002/cjp2.100.

[9] Ibrahim N, Lazaris A, Rada M, Petrillo S, Huck L, Hussain S, et al. Angiopoietin1 Deficiency in Hepatocytes A ff ects the Growth of Colorectal Cancer Liver. Cancers (Basel) 2020;12:1–18.

[10] Palmieri V, Lazaris A, Mayer T, Petrillo S, Alamri H, Rada M, et al. Neutrophils expressing lysyl oxidase–like 4 protein are present in colorectal cancer Therapy, metastases resistant to anti-angiogenic. J Pathol 2020. https://doi.org/10.1002/path.5449.

[11] Rada M, Lazaris A, Kapelanski-Lamoureux A, Mayer TZ, Metrakos P. Tumor microenvironment conditions that favor vessel co-option in colorectal cancer liver metastases: A theoretical model. Semin Cancer Biol 2020:1–13. https://doi.org/10.1016/j.semcancer.2020.09.001.

[12] Elmore S. Apoptosis: A Review of Programmed Cell Death. Toxicol Pathol 2007;35:495–516. https://doi.org/10.1080/01926230701320337.

[13] Chaitanya GV, Alexander JS, Babu PP. PARP-1 cleavage fragments: Signatures of cell-death proteases in neurodegeneration. Cell Commun Signal 2010;8:31. https://doi.org/10.1186/1478-811X-8-31.

[14] Rada M, Vasileva E, Lezina L, Marouco D, Antonov AV, MacIp S, et al. Human EHMT2/G9a activates p53 through methylation-independent mechanism. Oncogene 2017;36. https://doi.org/10.1038/onc.2016.258.

[15] Althubiti M, Rada M, Samuel J, Escorsa JM, Najeeb H, Lee KG, et al. BTK modulates p53 activity to enhance apoptotic and senescent responses. Cancer Res 2016;76:5405–14. https://doi.org/10.1158/0008-5472.CAN-16-0690.

[16] Nallanthighal S, Rada M, Heiserman JP, Cha J, Sage J, Zhou B, et al. Inhibition of collagen XI alpha 1-induced fatty acid oxidation triggers apoptotic cell death in cisplatin-resistant ovarian cancer. Cell Death Dis 2020;11:1–12. https://doi.org/10.1038/s41419-020-2442-z.

[17] Rada M, Barlev N, Macip S. BTK modulates p73 activity to induce apoptosis independently of p53. Cell Death Discov 2018;4:0–5. https://doi.org/10.1038/s41420-018-0097-7.

[18] Rada M, Barlev N, Macip S. BTK: a two-faced effector in cancer and tumour suppression. Cell Death Dis 2018;9:10–2. https://doi.org/10.1038/s41419-018-1122-8.

[19] Rada M, Nallanthighal S, Cha J, Ryan K, Sage J, Eldred C, et al. Inhibitor of apoptosis proteins (IAPs) mediate collagen type XI alpha 1-driven cisplatin resistance in ovarian cancer. Oncogene 2018. https://doi.org/10.1038/s41388-018-0297-x.

[20] Hu J, Bianchi F, Ferguson M, Cesario A, Margaritora S, Granone P, et al. Gene expression signature for angiogenic and nonangiogenic non-small-cell lung cancer. Oncogene 2005;24:1212–9. https://doi.org/10.1038/sj.onc.1208242.

[21] Coelho AL, Gomes MP, Catarino RJ, Rolfo C, Lopes AM, Medeiros RM, et al. Angiogenesis in NSCLC: Is vessel co-option the trunk that sustains the branches? Oncotarget 2017;8:39795–804. https://doi.org/10.18632/oncotarget.7794.

[22] Althubiti M, Rada M, Samuel J, Escorsa JM, Najeeb H, Lee K-G, et al. BTK modulates p53 activity to enhance apoptotic and senescent responses. Cancer Res 2016;76. https://doi.org/10.1158/0008-5472.CAN-16-0690.

[23] Rada M, Althubiti M, Ekpenyong-Akiba AE, Lee K-G, Lam KP, Fedorova O, et al. BTK blocks the inhibitory effects of MDM2 on p53 activity. Oncotarget 2017;8:106639–47. https://doi.org/10.18632/oncotarget.22543.

[24] Schippers IJ, Moshage H, Roelofsen H, Mu M, Ruiters M, Kuipers F, et al. Immortalized human hepatocytes as a tool for the study of hepatocytic. Cell Biol Toxicol 1997;13:375–86.

[25] Hang H, Liu X, Wang H, Xu N, Bian J, Zhang J. Hepatocyte nuclear factor 4A improves hepatic di ff erentiation of immortalized adult human hepatocytes and improves liver function and survival. Exp Cell Res 2017;360:81–93. https://doi.org/10.1016/j.yexcr.2017.08.020.

[26] Farra R, Dapas B, Pozzato G, Scaggiante B, Agostini F, Zennaro C, et al. Effects of E2F1 – cyclin E1 – E2 circuit down regulation in hepatocellular carcinoma. Dig Liver Dis 2011;43:1006–14. https://doi.org/10.1016/j.dld.2011.07.007.

[27] Holmgren L, O’Reilly M, Folkman J. Dormancy of micrometastases: balanced proliferation and apoptosis in the presence of angiogenesis suppression. Nat Med 1995;1:149–153.

[28] Valiente M, Obenauf AC, Jin X, Chen Q, Zhang XH, Lee DJ, et al. Serpins Promote Cancer Cell Survival and Vascular Co-Option in Brain Metastasis. Cell 2014;156:1002–16. https://doi.org/10.1016/j.cell.2014.01.040.

[29] Valiente M, Obenauf AC, Jin X, Chen Q, Zhang XH-F, Lee DJ, et al. Serpins Promote Cancer Cell Survival and Vascular Cooption in Brain Metastasis. Cell 2014;156:1002–16. https://doi.org/10.1124/dmd.107.016501.CYP3A4-Mediated.

[30] Donnem T, Reynolds AR, Kuczynski EA, Gatter K, Vermeulen PB, Kerbel RS, et al. Non-angiogenic tumours and their influence on cancer biology. Nat Rev Cancer 2018. https://doi.org/10.1038/nrc.2018.14.

[31] Brostjan C, Oehler R. The role of neutrophil death in chronic inflammation and cancer. Cell Death Discov 2020;6. https://doi.org/10.1038/s41420-020-0255-6.

